# Eurycomanone regulates lipid metabolism by activating the cAMP/PKA pathway

**DOI:** 10.1101/2020.08.20.258855

**Authors:** Zhihui Jiang, Haote Han, Shouxin Li, Jingkui Tian, Zhiyuan Gao, Wenping Huang, Dan Zhang, Hui Ouyang, Yulin Feng

## Abstract

*Eurycoma longifolia Jack* (ELJ) contains mainly alkaloids, and quassinoids, which are the main active ingredients. Eurycomanone (EN), one of the most common quassinoids, is said to have beneficial effects on lipid and glucose metabolism. In this study, we investigated the effects of EN on lipolysis by establishing a high-fat animal model *in vivo* and evaluated its efficacy as a lipolytic and anti-fatty liver agent. Oil red O staining showed morphological changes of 3T3-L1 preadipocytes after EN treatment and confirmed the inhibitory effects of EN on adipocyte differentiation. The mechanism of EN promotes lipolysis in 3T3-L1 cells was analyzed by immunofluorescence, Western blotting, quantitative real-time PCR and siRNA transfection. In C57BL/6J mice fed a high-fat diet, intragastric administration of EN (5 mg/kg and 10 mg/kg) for two weeks, decreased fat droplet mass and size, and reduced fat accumulation in the liver. Furthermore, EN activated PKA and promoted the PKA/hormone sensitive lipase lipolysis signaling pathway, thereby increasing the release of glycerol and free fatty acids from adipocytes. Our findings indicate the potential of EN as a promising alternative pharmacologic agent for the prevention of obesity.

## 1. Introduction

Obesity is a complex disease mainly caused by excessive accumulation of fat. When there is an imbalance between energy acquisition and energy consumption, the number of adipocytes gradually increases, the formation and storage of large lipid droplets. The incidence of obesity is increasing worldwide at an alarming pace and has become a major threat to public health [1]. Obesity is closely associated with type 2 diabetes, cardiovascular disease, non-alcoholic fatty liver disease and metabolic syndrome; therefore, the search for effective and safe weight-loss drugs has become a common aspiration.

Energy storage in the form of lipids in white adipose tissue (WAT) leads to WAT enlargement and eventually, to body weight gain [2–6]. Promoting WAT decomposition and inhibiting lipid formation can prevent excess lipid accumulation. Inhibit the differentiation of preadipocytes and promote lipolysis are proposed to be an alternative strategy to treat obesity [7].

Lipolysis is a complex process that is highly regulated and involved in coordinating several lipases and lipid droplet (LD)-associated proteins [8]. Adipose triglyceride lipase (ATGL), hormone sensitive lipase (HSL), and monoacylglycerol lipase are the three major lipases involved in lipolysis [9]. The activity of ATGL, which is highly specific for triacyl substrates, is largely determined by its comparative gene co-activation of 58 (CGI-58), whereas G (0)/G (1) switching gene 2 (G0S2) acts as an inhibitor of ATGL activity and ATGL-mediated lipolysis [10]. It has recently been shown that ATGL is phosphorylated at Ser406 by modulation of AMPK, leading to increased TAG hydrolase activity [11]. The activity of HSL is regulated by reversible post-transcriptional phosphorylation. In mouse adipocytes, PKA phosphorylates HSL at residues Ser563, 659 and 660, resulting in increased HSL translocation from the cytoplasm to the lipid droplet and lipase activity at the droplet surface [12]. In addition, AMP-activated protein kinase (AMPK) phosphorylates HSL at Ser565, thereby preventing PKA-induced HSL phosphorylation [13–14]. Activation of phosphodiesterase 3B (PDE3B) attenuates PKA activity by Akt-mediated phosphorylation of Ser273, thereby reducing HSL activation and lipolysis [15–16]. In addition to PKA-mediated phosphorylation, HSL can also be phosphorylated by other kinases, such as the extracellular signal-regulated kinase (ERK11\2), which activates HSL by phosphorylation at Ser600 [17]. Studies have also indicated that c-Jun-N-terminal kinase (JNK) plays a role in the regulation of lipolysis based on the silencing effect of Jnk1 and Jnk2 to promote lipolysis in mouse adipocytes [18]. Perilipin A forms a scaffold for lipid droplets, and plays an important role in protein coordination during lipolysis [19]. Under certain conditions, perilipin A restricts translocation of the lipase from the cytoplasm into the lipid droplets (LD), thereby reducing the rate of fat decomposition. However, PKA promotes perilipin A phosphorylation and conformational changes, which promotes the translocation of phosphorylated HSL to LDs, thereby increasing lipolysis [20]. Studies have indicated that a newly identified intracellular fat-specific phospholipase A2 (AdPLA) interferes with lipolysis by regulating the production of arachidonic acid [21, 22].

Eurycomanone (EN) is a naturally occurring quassinoids derived from *Eurycoma longifolia Jack* (ELJ) [Family: Simaroubaceae] (Figure 1). EN has been shown to exert a variety of physiological activities including anti-cancer effects, exhibiting strong dose-dependent anticancer efficacy against lung carcinoma (A-549 cells) and breast cancer (MCF-7 cells), while moderate efficacy was achieved against gastric (MGC-803 cells) and intestinal carcinomas (HT-29 cells) [23–25]. The pharmacokinetic properties of EN after oral administration of the pure compound and of *E. longifolia* extracts remain to be determined. The present method would be useful for future pharmacokinetic studies of the efficacy and safety of EN [26–27], metabolite research *in vivo* [28], toxicology [29–30], lipolysis and improvement of hypertension [31–33]. At present, there are a lot of theories related to lipolysis that have been supported in a large number of animal studies. The most common theory related to lipolysis involves the promotion of male sexual function. Sperm survival is associated with lower cholesterol levels in the testes. Increased lipolysis makes sperm more energetic, and shows synergy with other bioactive compounds that reduce body weight [34–35].

**Figure 1.**
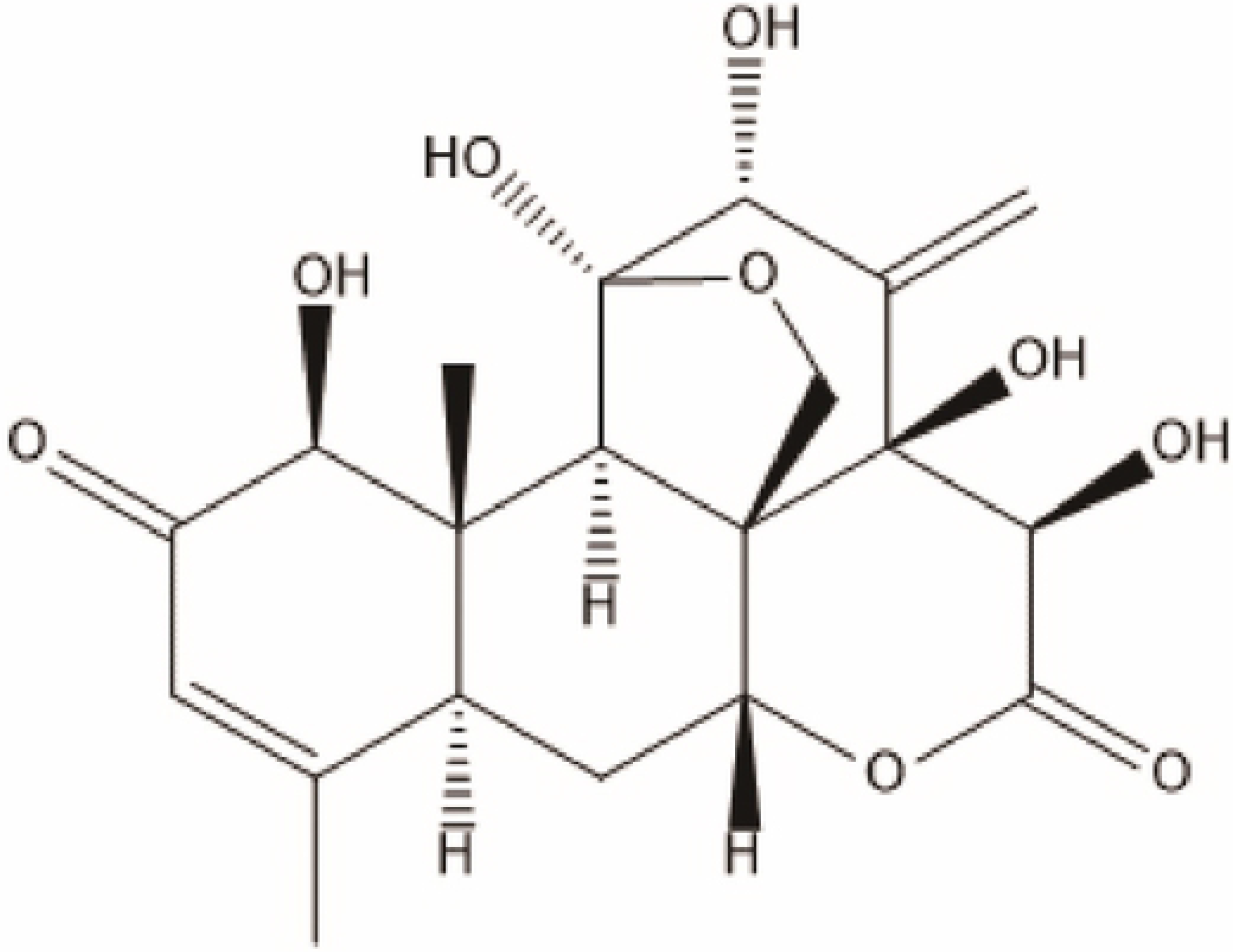
Chemical structure of Eurycomanone

In this study, we investigated the inhibitory effect of EN on adipocyte differentiation and lipid accumulation in 3T3-L1 preadipocytes, and assessed the expression level of genes contributing to lipolysis. We found that EN promotes AKT phosphorylation in 3T3-L1 adipocytes and stimulates lipolysis by activation of the cAMP/PKA pathway.

## 2. Materials and methods

### Reagents

The following monoclonal antibodies were used in this study: mouse anti-β-actin, rabbit anti-phosphorylated p-PKA, p-HSL, AKT and p-AKT (Cell Signaling Technology, 32 Tozer Road Beverly, MA, 01915 United States, 9621, 4126, 4691, and 4060), PKA, HSL, ATGL, cAMP (Abcam, ab75991, ab45422, ab109251, ab76238), anti-IgG-horseradish peroxidase (HRP), HRP-conjugated goat anti-rabbit IgG (H+L) and HRP-conjugated goat anti-mouse IgG-HRP (ImmunoResearch Laboratories) and FITC-conjugated goat anti-rabbit IgG (Beyotime, A0562). EN (purity ≥98%) was isolated by Jingkui Tian’s lab 49, the Key Laboratory of Biomedical Engineering, Zhejiang University China. High-glucose Dulbecco’s modified Eagle’s medium (DMEM), PKA inhibitor H-89 (S1643), insulin, dexamethasone (DEX), 3-(4,5-dimethyl-thiazol)-2,5-diphenyltetrazolium bromide (MTT), and 3-isobutyl-1-methylxanthine (IBMX) were purchased from Sigma–Aldrich; glycerol release assay kits were obtained from Nanjing Built Biotechnology Co., Ltd. Hematoxylin (C0107), eosin (C0109), DAPI (C1002), sodium citrate buffer (M019) and BSA (Bovine serum albumin, ST023) were from Gefan. Certified fetal bovine serum (FBS) was from BI. Methanol, ethanol, xylene, dimethyl sulfoxide (DMSO), oil red O and formaldehyde solution were from Sinopharm, Nero red were from Huaan Biotechnology Co., Ltd., China.

### Animals and treatment

Male C57BL/6 mice (aged 12 weeks, batch SCXK (Su) 2018-0008) were acclimatized with 12 h dark-light cycles under a constant temperature (22 ± 2°C) and were given free access to food and water. All experiments were carried out in accordance with the internationally accepted guidelines for the care and use of laboratory animals and were approved by Animal Ethics Committee of School of Chinese Materia Medica, Jiangxi University of Traditional Medicine (Nanchang, China).

The high-fat animal model was established by feeding mice for 8 weeks with a high-fat diet (HFD, 10% lard, 10% yolk, 1% cholesterol, 0.2% cholate and 78.8% standard diet, Nanjing Qinglongshan Experimental Animal Center). Mice received EN (5 mg kg^-1^, 10 mg kg^-1^) by intragastric administration. Two weeks later, animals were fasted for 24 h before blood samples were collected from the orbital sinus and glycerol levels were assayed with commercial kits. Mice were sacrificed by cervical dislocation and epididymal adipose tissues and liver were isolated and stored at −80°C for assay. Other parts of liver as well as epididymal and perirenal white adipose tissues were excised, weighed and portions of the tissues were fixed in 10% formalin for histopathology.

### Hematoxylin and eosin (H&E) staining

Tissues were dissected and fixed overnight in 4% paraformaldehyde. Paraffin processing, embedding, sectioning, and standard H&E staining was performed. Images were acquired using an inverted microscope (Nikon).

### Cell culture and differentiation

Murine 3T3-L1 preadipocyte (ATCC® CL-173™) cells were cultured in DMEM with 10% FBS (10% FBS/DMEM) and antibiotics (100 units/mL penicillin, 100 μg/mL streptomycin) maintained at 37°C under a humidified atmosphere of 5% CO_2_ in an incubator. Adipocyte differentiation was induced one day after reaching confluence (day 0) by changing the medium to DMEM with 10% FBS and 0.5 mM IBMX, 0.25 μM DEX and 10 μg/mL insulin (differentiation medium). Two days after induction (day 3), the medium was replaced with DMEM with 10% FBS and 10 μg/mL insulin medium to enhance differentiation, and the cells were cultured for another 2 days. On day 5, the medium was replaced with DMEM with 10% FBS and 10 μg /mL insulin medium and the cells were cultured for a further 2 days cells. Then cells were cultured with DMEM with 10% FBS medium for two more days. On day 8, cells were used in glycerol release enhancement assays.

### Cell viability assay

Cell viability was determined by 3-(4, 5-dimethylthiazol-2-yl)-2,5-diphenyltetrazolium bromide (MTT) assay. 3T3-LI preadipocyte cells were seeded in 96-well plates at 5000 cells/well (n = 6 replicates) in a final volume of 100 μL. After incubation for 24 h at 37°C under 5% CO_2_, cells were treated with EN (at final concentrations of 0, 25, 50, 100, or 200 μM). After incubation for 24 h and 48 h, cells were treated with 50 μL of the MTT (5 mg/mL stock solution) per well for another 2 h. The medium was then carefully removed from each well and replaced with 200 μL DMSO. Absorbance was measured at 570 nm on a plate reader (81 Wyman Street, Waltham, MA, 02454, USA).

### Oil red O staining

3T3-L1 cells were washed twice with PBS and then fixed with 10% formaldehyde for 2 h at room temperature. The formaldehyde was then discarded and cells were rinsed with 60% isopropanol for 5 min before staining for 1 h with oil red O working solution (60% oil red O stock solution: 40% deionized water). The stained cells were immediately washed with deionized water and then photographed under a microscope.

### Glycerol release assay

On day 8, the medium was changed to sample-containing medium (phenol-red-free DMEM) and incubated for 6 h. In inhibitor studies, the cells were incubated with inhibitor for 1 h prior to addition of the EN, and then incubated for a further 24 h. On the day of the glycerol release enhancement assay, the medium was collected and mixed with a free glycerol reagent (Glycerol release assay kit, Nanjing Built Biotechnology Co., Ltd. China). The mixture was incubated at 37°C for 5 min and its absorbance at 550 nm was measured to quantify the amount of the released glycerol. The absorbance relative to that of the negative control was calculated. Isoproterenol hydrochloride 1 μM (Sigma–Aldrich Co., St Louis, MO, USA) was used as positive control.

### Western blotting analysis

Cells seeded in 6-well plates (5×10^5^/well) were treated with isoproterenol (ISO) or pre-treated with H-89 (Inhibits. PKA inhibitor, final concentration 5 μg/mL) for 1 h and then treated with EN at various concentrations for 24 h. Cells were then collected from 6 cm dishes by scraping and centrifugation at 500 *×g* for 5min. The collected cells were then washed with ice-cold PBS, resuspended in 60–100 μL lysis buffer and sonicated for 15–20s. Lysates were centrifuged at 12000 *×g* for 15 min at 4°C. Protein concentrations in the supernatants were determined by using a BCA (3rd Floor, 85A, U-Valley, Liandong, Tongzhou District, Beijing, China) protein concentration determination kit. Equal amounts of proteins were separated by SDS polyacrylamide gel (7.5%) electrophoresis and then transferred to a polyvinylidene fluoride (PVDF) membrane. The membrane was blocked with 5% non-fat milk in 0.1% Tween-20 in TBS (TBST) for 2 h and incubated with primary detection antibody (1:500–1:1000) at 4°C overnight. After being washed three times with TBST, the membrane was incubated for 2 h at room temperature with the appropriate secondary antibody (1:2000). The immunoblots were visualized with an enhanced chemiluminescence (ECL, Biyuntian Biotechnology, Building 30, Songjiang Science and Technology Entrepreneurship Center, Lane 1500, Xinfei Road, Songjiang District, Shanghai, China) system.

### Immunofluorescence analysis

Following treatment with EN, cells were washed with cold PBS, fixed with 4% paraformaldehyde (chilled to −20°C) for 20 min at room temperature, permeabilized with 0.2% Triton X-100 in PBS for 5 min, and then blocked with 5% BSA in PBS for 30 min. After being washed with PBS, the cells were incubated overnight with P-HSL antibody in 1% BSA at 4°C. The cells were then incubated with FITC goat anti-rabbit IgG in 1% BSA for 2 h at room temperature. Nuclei were stained with DAPI. Confocal images were obtained under oil immersion lens using a confocal microscope (LSM 780, Carl Zeiss Jena, Germany).

### RNA isolation and quantitative real-time PCR (qRT-PCR)

Total cellular RNAs were extracted using the TRIzol reagent (Invitrogen) according to the manufacturer’s instructions. cDNA was synthesized from the isolated RNA by reverse transcription using RevertAid™ first strand cDNA synthesis kit (5× All-In-One RT MasterMix, ABM) according to the manufacturer’s instructions in a RNase-free environment. The relative expression of genes was quantified by quantitative real-time polymerase chain reaction (qRT-PCR) analysis using SYBR Green Supermix (TaKaRa); β-actin served as a housekeeping gene. qRT-PCR was performed using the CFX Connect system (Bio-Rad), and the relative expression of genes was analyzed using the 2^−ΔΔCT^ method. Three independent experiments were performed, and the primer sequences are listed in Table 1.

**Table. 1.**
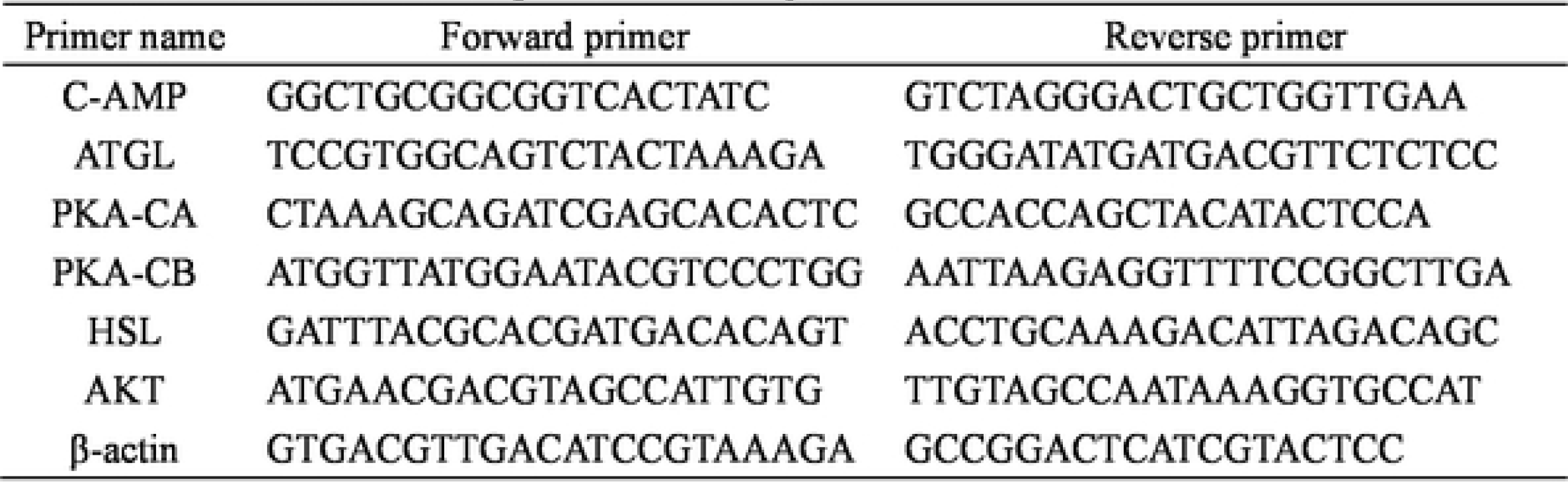
Primers used for qRT-PCR analysis.

### PKA-ca siRNA transfection

Cells were seeded in 6-well plates (5×10^5^/well) were transfected with the indicated siRNA (3 μg) using Lipofectamine® 2000 transfection reagent (Life Technologies, 11668-019) according to the manufacturer’s instructions. Treatments were incubated for 48 h after transfection 24 h. The siRNA-PKA-ca was purchased from Shangya Biotechnology Co., Ltd (Zhejiang, china).

### Statistical analysis

Data were expressed as means ± standard deviation (SD). Data were analyzed by using analysis SPSS software (SPSS Inc., Chicago). The significance of the difference among mean values was determined by one-way analysis of variance followed by Tukey’s test. Comparisons between two groups were made using Student’s *t*-test. A value of *P* < 0.05 was considered to indicate statistical significance.

## 3. Results

### 3.1 EN cytotoxicity in 3T3-L1 adipocytes

The cytotoxicity of EN on preadipocytes and mature adipocytes were evaluated using MTT and Trypan blue assays. The MTT assay indicated that culturing cells with EN had no adverse effects on preadipocyte viability. The Trypan blue assay also indicated that treating mature adipocytes with EN had no adverse effects on cell viability (Fig. 2A and B, respectively).

**Figure 2.**
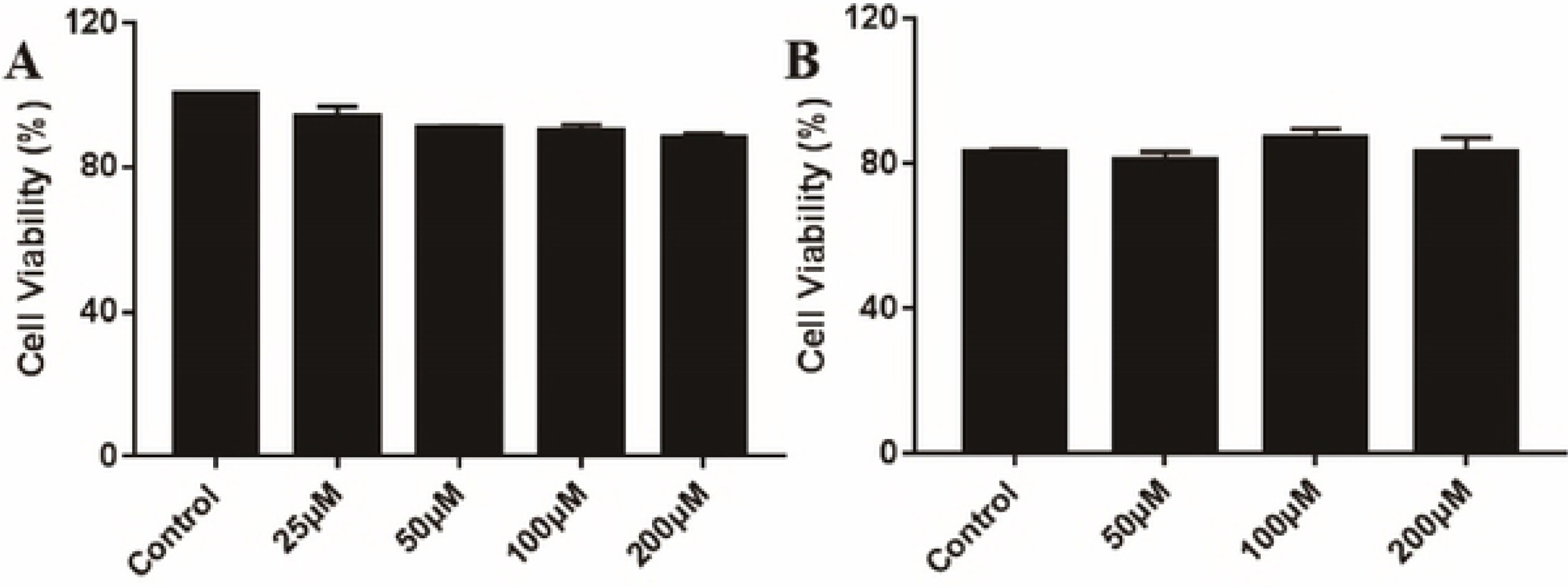
3T3-L1 pre-adipocytcs were treated with 25, 50,100 and 200 EN 72 h. (A) 3T3-L1 adipocytes were treated with 50,100 and 200 EN 24 h (B). The level in control cells was regarded as 1. Cell viability was assessed by MTT assay. The level in control cells was regarded as 100%. Values arc means ± SD (n=4). According to the student’s t test, there was no significant difference between the dosing group and the control cells.

### 3.2 EN inhibited lipid accumulation and reduced TG contents in differentiating 3T3-L1 cells

Lipid accumulation is the most prominent marker of adipogenesis, and its quantification is used to assess the extent of adipocyte differentiation. Intracellular fat accumulation was significantly decreased by treatment with EN for 4 days compared with control. The lipid content in EN-treated cells during the early phase of adipocyte differentiation was lower than that in intermediate and late stages (Fig. 3A and B). These results suggested that EN influenced the early phase of adipogenic differentiation. Several concentrations of EN (15 μM, 30 μM and 45 μM) were tested to investigate its inhibitory effect on adipocyte differentiation, simultaneously verifying the period of EN inhibition of cell differentiation. EN was added to 3T3-L1 cells together with 3-isobutyl-1-methylxanthine, dexamethasone, and insulin (MDI) stimulation, the degree of differentiation was quantified according to LD accumulation quantified by oil red O staining and observed by microscopy (Fig. 3C and D). EN at 15 μM, 30 μM and 45 μM inhibited adipocyte differentiation by 39.7%, 45.6% and 44.4% respectively. These effects were further confirmed by Nero red staining which observed by laser confocal imaging (Fig. 3E). Adipocyte differentiation comprises early, intermediate and late phases. To identify the crucial stage of adipocyte differentiation influenced by EN treatment, 3T3-L1 preadipocytes were treated with EN (30 μM) at various times during adipogenesis, and intracellular fatty accumulation was measured by quantitative analysis of oil red O-stained intracellular LDs. We then investigated the lipolytic effects EN in 3T3-L1 adipocytes by measurement of glycerol contents in the culture medium of EN-treated cells. Triglycerides (TGs) are hydrolyzed to form free fatty acids and glycerol in the event of energy demand. Glycerol content hence indicates the degree of adipolysis in mature adipocytes. Treatment of mature adipocytes with EN (30 μM) for 24 h significantly induced glycerol release (35%) (Fig. 3F).

**Figure.3.**
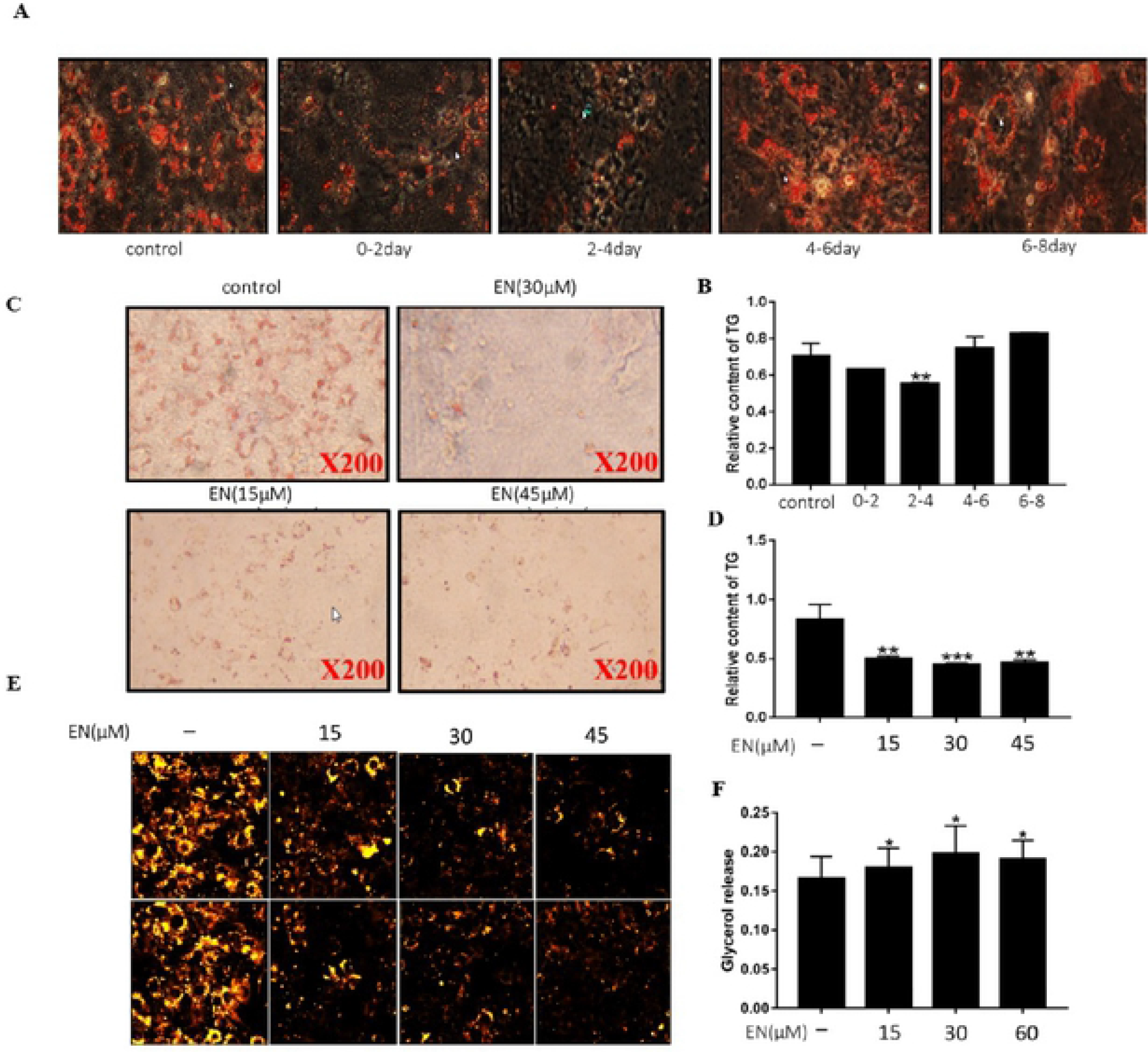
(A) shows the results of adding EN (30 μM) at four different time periods of 0-2 days, 2-4 days, 4-6 days, 6-B days. (B) Accumulation of intracellular triglycerides after addition of 3T?1 prc-adipocytes with EN at different time points. (C) Cell differentiation morphology of 3T3-L1 prcadipocytcs at 15 μM, 30 μM, 45 μM and 2-4 days after treatment with EN. (D) Accumulation of intracellular triglycerides after 3T-?1 prcadipocytcs were treated with EN. (E) Cell differentiation morphology of adipocytes under laser confocal microscopy at !5μM, 30 μM, and 45μM. (F) Results of glycerol release under the action of three concentrations of EN at 15 μM, 30 μM, and 45 μM. Values arc means ± SO (n=3). Data were analyzed by using Student’s t-test. *p < 0.05, **p < 0.01, ***p < 0.005 Compared with the control.

### 3.3 EN promotes adipolysis via the cAMP/PKA/HSL signaling pathway

Western blotting analysis was conducted to determine the effect of EN on cellular protein levels of cAMP, p-PKA and p-HSL after treatment for 24 h. MDI stimulation enhanced TG accumulation in 3T3-L1 adipocytes; however, EN treatment increased protein expression levels of p-PKA and p-HSL, and promoted TG decomposition (Fig. 4A, B). These results were consistent with qRT-PCR analysis of these genes (Fig. 4C). ISO, a well-known lipolytic agent, was used as a positive control. In differentiated 3T3-L1 adipocytes, ISO stimulated higher levels of glycerol release than that stimulated by EN. Moreover, both ISO and EN strongly induced PKA and HSL phosphorylation, but had no significant effects on the level of total HSL protein (Fig. 4D). The densitometry data are shown in Figure 4E, F. The Western blotting data were consistent with qRT-PCR analysis of these genes (Fig. 4G). We further investigated the role of the PKA/HSL signaling pathway in the effects of EN on adipocytes by using H-89 to inhibit PKA protein expression. H-89 treatment decreased PKA phosphorylation and HSL protein expression. EN treatment for 24 h significantly enhanced PKA phosphorylation regardless the presence or absence of H-89 *(P* < 0.05, Fig. 4H); this was consistent with the qRT-PCR analysis of these genes (Fig. 4I). The Western blotting results were confirmed by immunofluorescence analysis, which showed that EN treatment promoted p-HSL protein expression in adipocytes (Fig. 4J).

**Figure.4.**
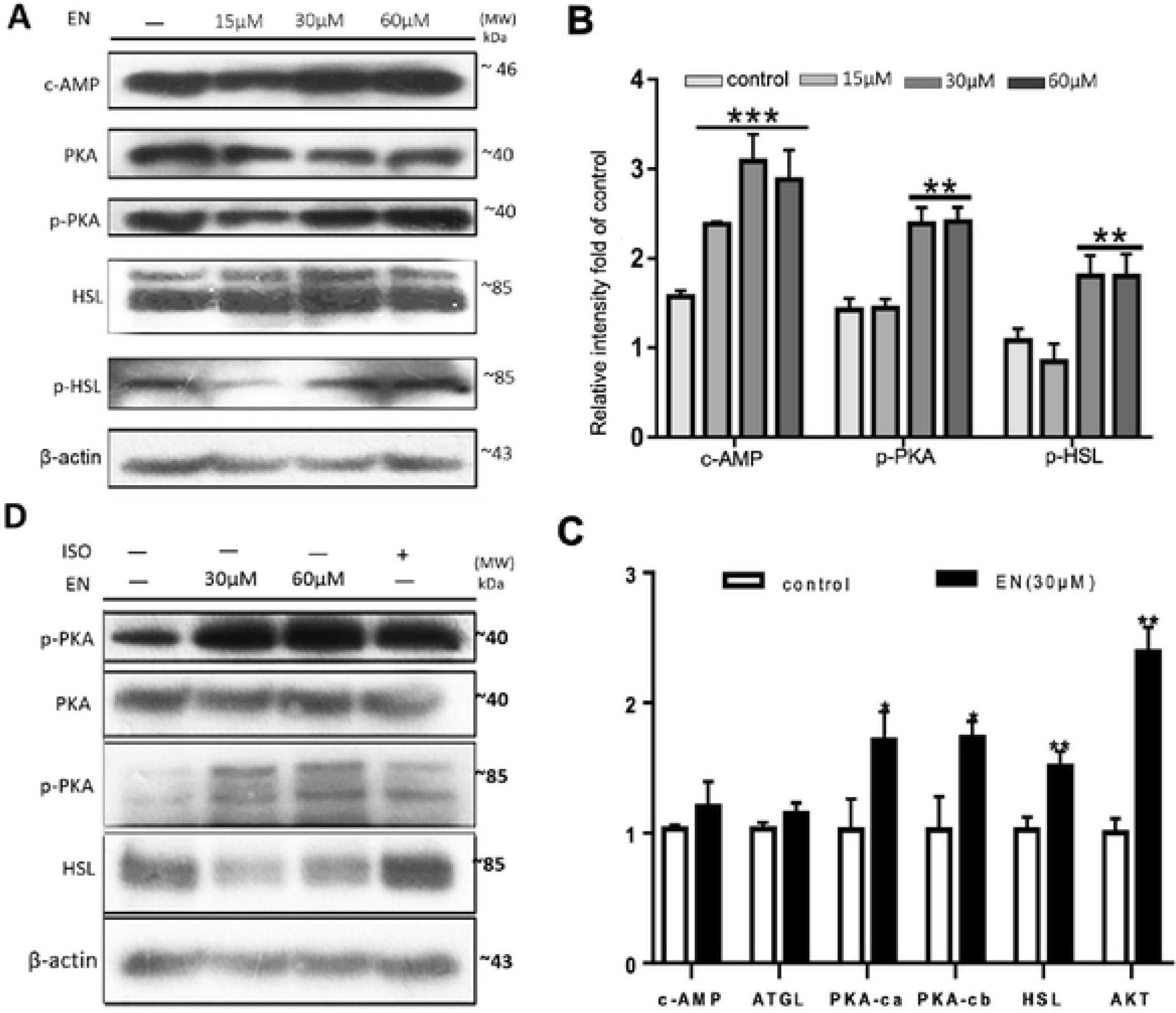
3T3-L1 adipocytes were pretreated with 30 μM EN for 24 h. Protein expression was determined by Western blot assay. One representative blot from three independent experiments is shown. The protein levels of the bands were quantified by densitometry. The effect of protein levels on C-AMP, P-PKA and P-HSL and the gray value after 24 hours of treatment of cells, (C-D) indicates that EN treated C-AMP, ATGL, PKA-CA, PKA-after 24 hours of treatment of cells. The effect of PKA-CB, AKT and HSL mRNA expression levels. (EF) compared to the positive control group, the effect of EN on the protein levels of P-PKA and P-HSL after 24 hours of treatment, and the gray value, (G) The effect of EN on the mRNA expression levels of C-AMP, ATGL, PKA-CA, PKA-CB, AKT and HSL after 24 hours of treatment with positive control group, (H-I) indicates the effect of EN after the action of PKA inhibitor H89 And the effect of ISO on the protein levels of C-AMP, P-PKA and P-HSL after 24 hours of treatment of cells and the gray value. (J) P-HSL protein expression under laser confocal scanning microscopy. The level in control cells was set at 1. Values arc means ± SD (n=3), Data were analyzed by using Student’s t-test. *p < 0.05, **p < 0.01, ***p < 0.005 Compared with the control.

### 3.4 Effect of EN on AKT signaling pathways

The effect of EN on p-AKT protein level in cells was determined by Western blotting assay. After EN treatment, p-PKA expression was significantly increased, whereas glycerol release was decreased (Fig. 5A–D). In the presence of the AKT inhibitor (MK), glycerol release from adipocytes was increased. However, in the presence of both EN and MK were combined, glycerol release from adipocytes was significantly inhibited (Fig. 5E–F).

**Figure.5.**
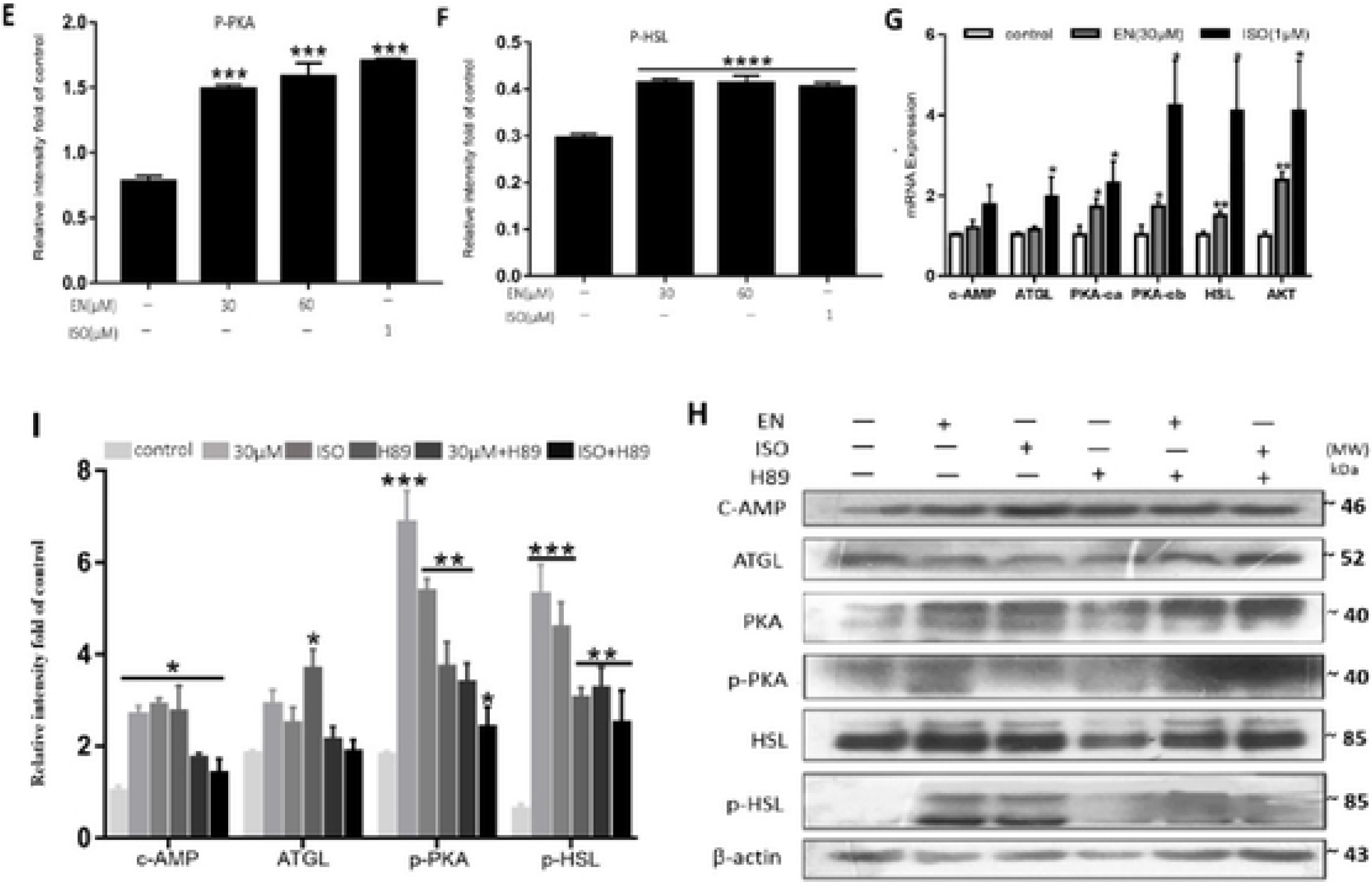
3T3-L1 adipocytes were pretreated with 30 μM EN for 24 h. Protein expression was determined by Western blot assay. (E-F) compared to the positive ∞ntrol group, the effect of EN on the protein levels of P-PKA and P-HSL after 24 hours of treatment, and the gray value, (G) The effect of EN on the mRNA expression levels of C-AMP, ATGL, PKA-CA, PKA-CB, AKT and HSL after 24 hours of treatment with positive ∞ntrol group, (H-I) indicates the effect of EN after the action of PKA inhibitor H89 And the effect of ISO on the protein levels of CAMP, P-PKA and P-HSL after 24 hours of treatment of cells and the gray value. Data were analyzed by using Student’s t-test. *p < 0.05, **p < 0.01, ***p < 0.005 Compared with the control.

**Figure.5-1.**
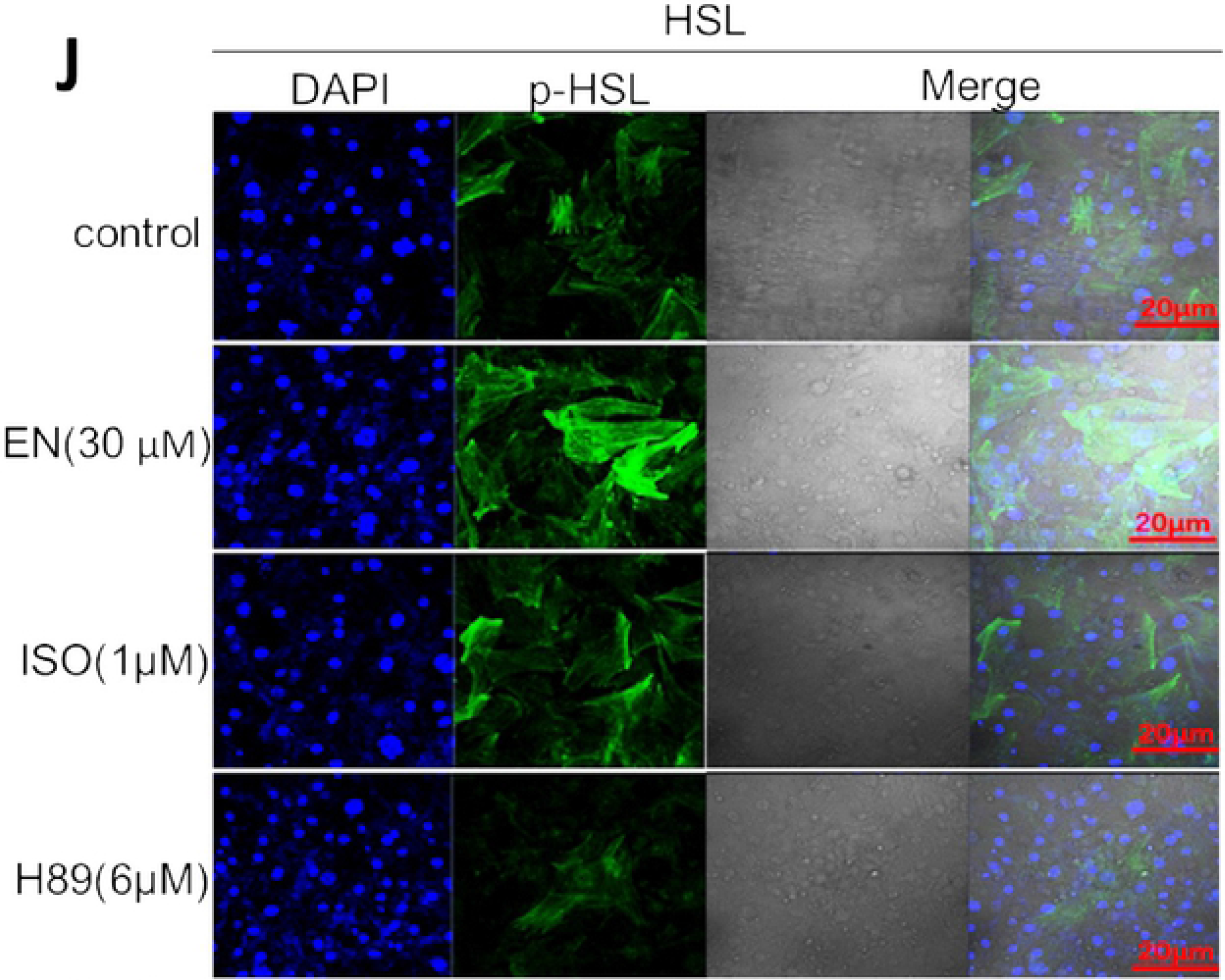
Following treatment with EN, The Western blotting results were confirmed by immunofluorescence analysis. (J) P-HSL protein expression under laser confocal scanning microscopy. The level in control cells was set at 1. Values are means ± SD (n=3). Data were analyzed by using Student’s t-test. *p < 0.05, **p < 0.01, ***p < 0.005 Compared with the control.

### 3.5 Effect of PKA-ca-silencing on lipolysis in 3T3-L1 cells

Compared with the untransfected control group, transfection of 3T3-L1 adipocytes with control siRNA had no significant effect on glycerol release (*P* > 0.05), whereas glycerol release was significantly increased in siRNA-mediated silencing of PKA-ca group after treatment with EN (*P* < 0.05). These results indicated that EN promotes the breakdown of intracellular TG (Fig. 6A). Western blot analysis showed that compared with the normal group, p-PKA protein expression was significantly inhibited by siRNA transfection (*P*<0.05), while p-HSL protein expression was significantly increased (*P* < 0.05). Immunofluorescence analysis showed that p-HSL protein expression was significantly increased compared with that in the control group (*P* < 0.05), while the expression of p-PKA protein was significantly increased (*P* < 0.05), indicating that PKA and HSL are involved in the regulation of lipolysis (Fig. 6B). Following siRNA-mediated silencing of PKA-ca, mRNA expression of cAMP, PKA-ca, PKA-cb and HSL in adipocytes treated with EN for 24 h was measured by qRT-PCR analysis. There was no significant change in PKA-cb expression, whereas PKA-ca expression was significantly inhibited. HSL expression was also inhibited. However, there was no significant change in PKA-cb expression following the addition of EN (30 μM), whereas the expression of PKA-ca and HSL was up-regulated (Fig. 6C).

**Figure.6.**
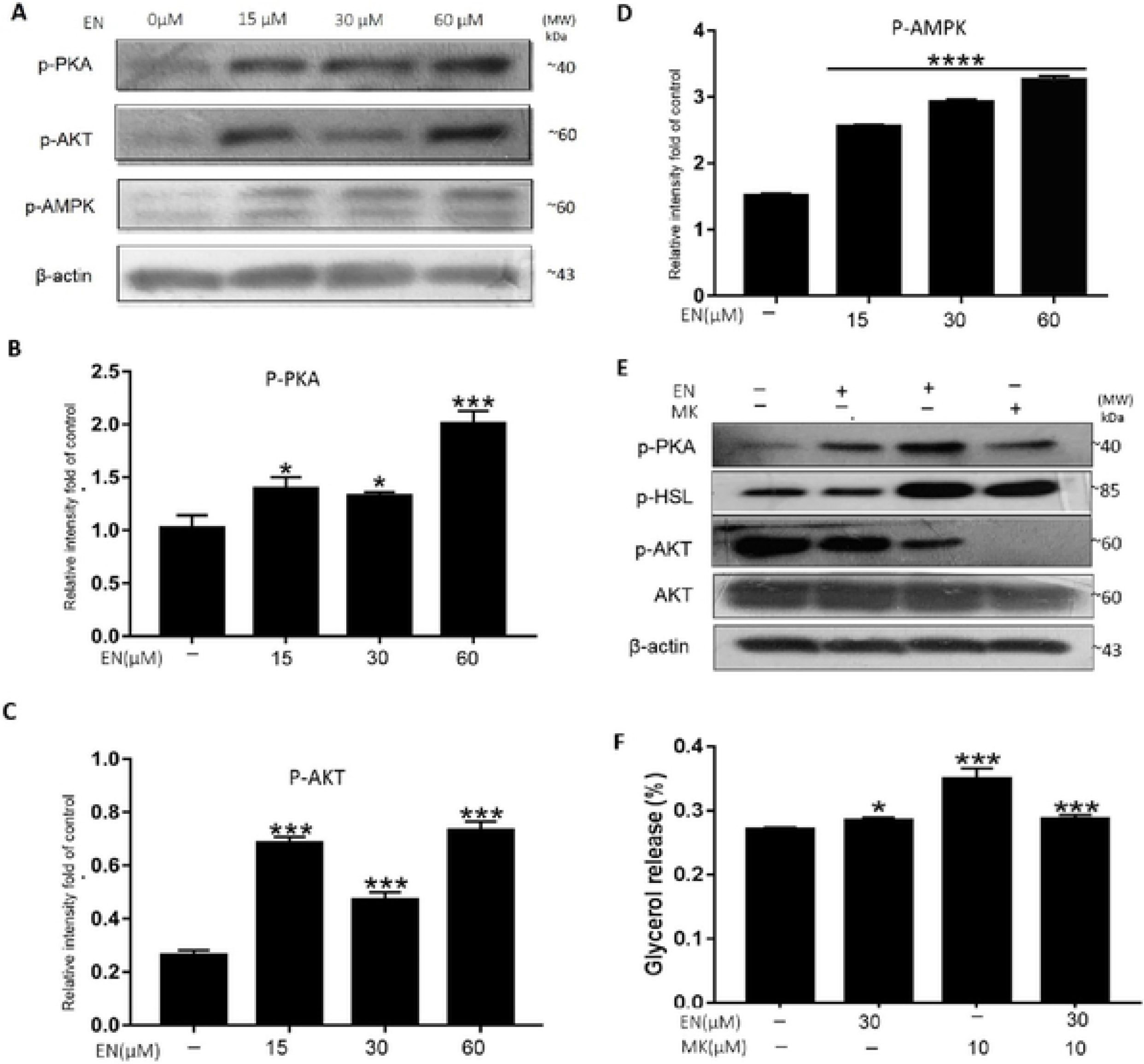
3T3-L1 adipocytes were pretreated with 30 μM EN for 24 h. Protein expression was determined by Western blot assay. One representative blot from three independent experiments is shown. The protein levels of the bands were quantified by densitometry. The effect of P-AKT, P-PKA and P-HSL on protein levels and gray-scale W values after 24 hours of treatment of cells, (E) indicates EN and MK versus P-AKT, P-PKA and P-HSL after 24 hours of treatment of adipocytes, Results of glycerol release under the action of concentrations of EN at 30 μM, MK. Values arc means ± SD (n=3). Data were analyzed by using Student’s t-test. * p < 0.05, **p < 0.01, ***p < 0.005 Compared with the control.

### 3.6 Effects of EN treatment on lipid profile in C57BL/6J mice

Serum TG levels were elevated in C57BL/6J mice fed HFD compared with control group. Intragastric administration of EN significantly decreased TG levels and increased serum glycerol release in mice fed HFD (Fig. 7A–B). Furthermore, H&E staining showed a reduced adipocyte size as well as a reduction in the number of LDs in the liver of C57BL/6J fed HFD following treatment with EN (Fig. 7C).

**Figure.7.**
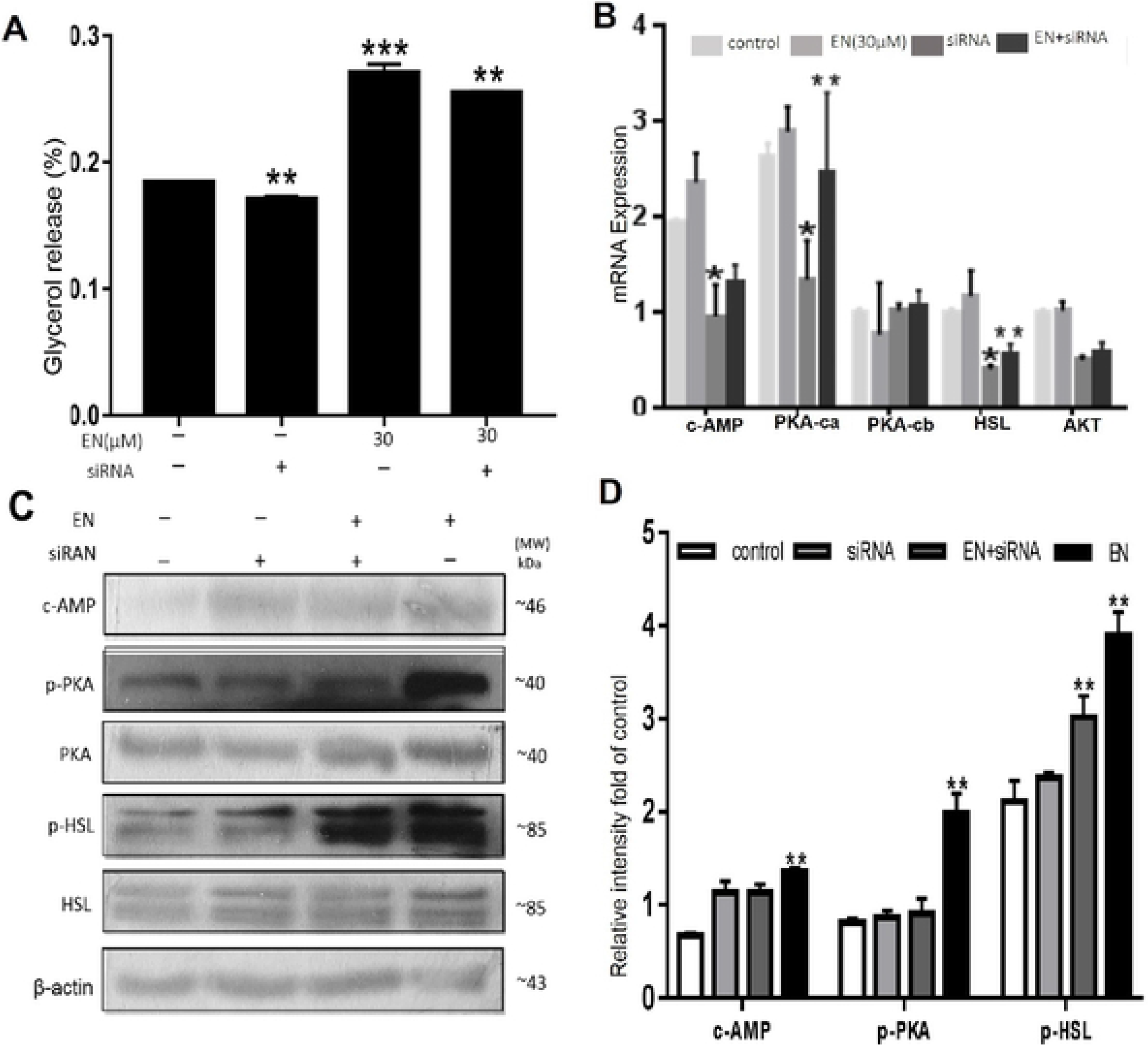
Silence PKA-CA Gene pair 3T3-L1 Effect of lipolysis. (A) shows the results of glycerol release after treatment of EN (30 μM), siRNA, EN (30 μM) + siRNA on adipocytes, (B) indicates EN (30 μM), siRNA The effect of protein levels on C-AMP, P-PKA and P-HSL (C) indicates that the treated after 24 hours of treatment of cells, the effect of C-, EN (30 μM) + siRNA AMP, PKACA, PKA-CB, AKT and HSL mRNA expression levels. The level in ∞ntrol cells was set at 1. Values arc means ± SD (n=3), Data were analyzed by using Student’s t-test. *p < 0.05, **p < 0.01, ***p < 0.005 Compared with the ∞ntrol.

### 3.7 EN promoted cAMP/PKA activation in adipose tissues

We validated the effects of EN on the cAMP/PKA pathway in HFD-fed mice. In accordance with the *in vitro* results, oral administration of EN promoted p-PKA induction in the adipose tissue of HFD-fed mice. In addition, PKA enzymatic activity assays showed that HFD feeding attenuated p-PKA activity in adipose tissue of HFD-fed mice, while EN administration promoted p-PKA activity (Fig. 8A–D).

**Figure.8.**
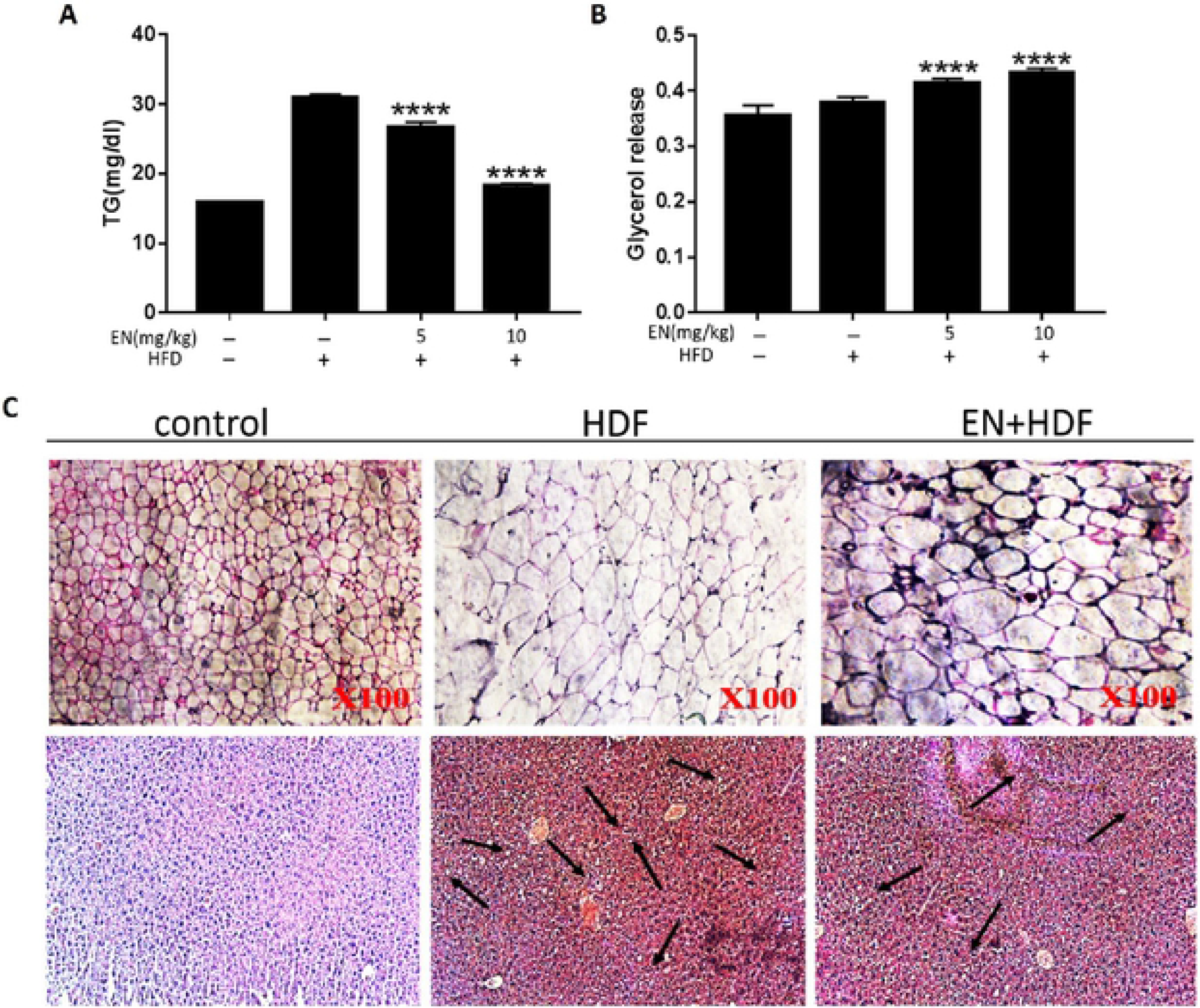
Effects of EN treatment on lipid profile in C57BL/6J mice. (A) Triglyceride content in mouse scrum, (B) glycerol ∞ntent in mouse scrum, (C) HE staining of liver tissue and adipose tissue. Values arc means ± SD (n=3), Data were analyzed by using Student’s t-test. ****p < 0.005 Compared with the control.

**Figure.9.**
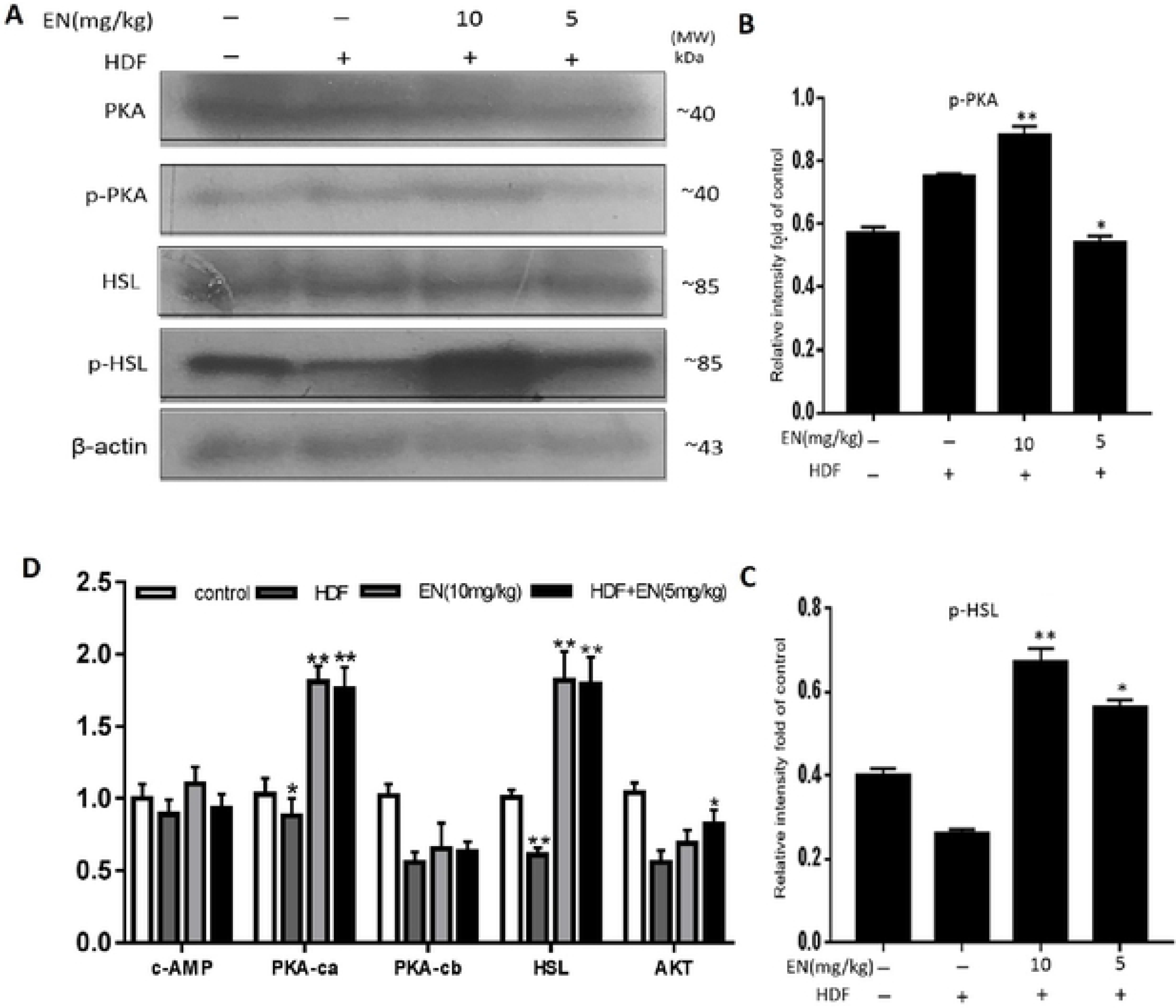
EN promoted cAMP/PKA activation in adipose tissues. (A-C) The effect of P-PKA and P-HSL on protein levels and gray-scale W values, (D) The effect of EN on the mRNA expression levels of C-AMP, PKA-CA, PKA-CB, HSL, AKT. Values arc means ± SD (n=3). Data were analyzed by using Student’s t-test. *p < 0.05, **p < 0.01, ***p < 0.005 Compared with the control.

## 4. Discussion

With the improvement in living standards, the incidence of obesity and related diseases has increased year by year, and the burden on families and society has also increased. Adipocyte metabolism and dysfunction play a pivotal role in obesity [36–37], which are currently focus of research. Abnormal expression of PKA phosphorylation has been identified in obese samples and closely related to lipid metabolism disorders [38–40]. The occurrence and development of obesity is inseparable from the accumulation level of TG, and the activation of protein kinases promotes TG decomposition in adipocytes. Adipocyte formation depends on pre-adipocyte differentiation. Thus, inhibiting the formation of adipocytes and promoting the decomposition of mature adipocytes is an effective method to induce weight loss. A recent study demonstrated that EN played a significant role in alleviating lipid accumulation and promoting lipolysis [41–42]. However, the role of EN in lipolysis and lipid accumulation is to be elucidated.

In this study, oil red O staining showed that adipocytes differentiated normally, whereas EN treatment resulted in smaller, misshapen LDs. Fat metabolism refers to the process of synthesis and decomposition of fat in living organisms. Both processes regulate fat accumulation and abnormal lipid metabolism leads to excessive lipid accumulation and obesity. In this study, TG accumulation in adipocytes was significantly lower following EN treatment compared with the levels detected in the normal group. Our results indicate that EN acts primarily in the early stages of adipocyte differentiation. *In vitro,* adipocytes are the basic tool for studying fat-related problems. The differentiation of adipocytes is an important part of the study of adipocytes-related problems. Adipocyte differentiation is accompanied by a change in morphology from the fusiform to the spherical shape, at which point, LDs begin to appear, indicating that the preadipocytes have differentiated into adipocytes. As the LDs accumulate, mature adipocytes are formed. Adipocyte differentiation has a very close relationship with energy balance, obesity, type 2 diabetes, fatty liver, hyperlipidemia and breast cancer [43–44]. The study of adipocyte differentiation is of great theoretical significance not only for the study of normal biological and disease processes, but also for the prevention and treatment of those diseases, and especially for screening the effects of drugs on cells and molecules. Therefore, comprehensive understandings of the mechanisms of adipocyte differentiation for reducing lipid levels are crucial to the prevention and treatment of these diseases. In this study, we established a model of adipocyte differentiation (≥ 90%) for the evaluation of the effects of EN on lipid metabolism.

In this study, we explored the effects of EN at the optimal dose of 30 μM on cAMP/PKA signal transduction in lipolysis in vitro at the protein and gene expression level by Western blotting, immunofluorescence, qRT-PCR, siRNA-mediated silencing and other techniques. Previous studies [45–48] have shown that the expression of PKA, HSL and cAMP mRNA is upregulated following EN treatment, while AKT expression is downregulated. However, in this study, we found that expression of PKA, HSL, cAMP and AKT were upregulated at both the mRNA and protein levels following EN treatment. Thus, our findings related to the expression of AKT following EN treatment are inconsistent with previous reports [49–52]. We speculated that the cAMP/PKA signaling pathway has a greater effect on lipolysis than the upregulation of AKT expression, such that EN promotes lipolysis. Our study also showed that EN had no effect on ATGL expression. PKA inhibition assays were conducted to verify that PKA is the major protein regulated by EN in promoting lipolysis. These studies confirmed that EN enhances the effects of H-89 inhibition. The pathway culminates in regulation of HSL, so we visually observed the effect of EN on the expression of HSL by immunofluorescence analysis. Our experimental results are consistent with previous reports [53–54]. Finally, we performed siRNA-mediated silencing of PKA to confirm its role in the mechanism by which EN promotes lipolysis. PKA-ca expression was significantly inhibited, and HSL expression was also inhibited. There was no significant change in PKA-cb expression following the addition of EN, while expression of PKA-ca and HSL was promoted. These results indicate that EN promotes lipolysis by acting on the PKA-ca site of PKA.

## 5. Conclusion

In this study, we showed that EN significantly attenuated lipid accumulation and differentiation in 3T3-L1 cells. Intervention with EN also promoted the expression of the main transcriptional regulator PKA and HSL followed by upregulation of adipogenic-specific molecules including CAMP, ATGL, PKA-CA, PKA-CB, HSL and AKT at the mRNA and protein levels, probably via the cAMP/PKA signaling pathway. In addition, the administration of EN decreased body weight gain and adipose tissue hypertrophy induced by HFD *in vivo*. Moreover, supplementation of EN improved glucose clearance and decreased TG levels in HFD-fed mice. Additionally, the development of hepatic steatosis was also significantly prevented in the obese mouse supplemented with EN. Collectively, our results strongly suggest the novel effect of EN in inhibiting adipogenesis, promoting lipolysis and the high therapeutic potential of EN in preventing the development of obesity.

## Acknowledgments

The study was funded by grants (No. 81560636, 81760702) from Natural Science Foundation of China, China Postdoctoral Science Foundation (No.2017M612159), JIANGXI Postdoctoral Science Foundation (No.2017KY07), Jiangxi Province 5511 innovative talent Project [20165BCB19009], Nan-chang innovative talent team (No. [2016]173).

